# Once in a blue moon: an unexpected evolutionary route to blue floral pigmentation in orchids

**DOI:** 10.64898/2026.10.20.760294

**Authors:** Emma Grace John, Ryan P. O’Donnell, James Perkins, Monica Rubial, Takayuki Mizuno, Rod Peakall, Darren C. J. Wong

**Affiliations:** Ecology and Evolution, Research School of Biology, The Australian National University, Canberra, ACT, 2600 Australia; Department of Botany, National Museum of Nature and Science, Tsukuba 305-0005, Japan; School of Agriculture, Food, and Wine, Waite Research Precinct, Adelaide University, South Australia, 5064, Australia

## Abstract

Blue flowers are rare in the hyperdiverse orchid family, and the genetic mechanisms enabling their blue delphinidin-based pigmentation remain poorly understood. Within the Australian orchid subtribe Caladeniinae (∼350 spp.), repeated transitions from maroon to blue flowers, followed by reversions to maroon, provide a unique opportunity to investigate blue flower evolution. Using phylotranscriptomics, anthocyanin profiling, molecular evolutionary analyses, and gene expression data from 51 species, we reconstructed the genetic changes associated with these transitions. Despite its near-universal role in delphinidin biosynthesis across flowering plants, unexpectedly, we found no evidence for a functional CYP75A-mediated flavonoid 3′5′-hydroxylase (F3′5′H) pathway. Instead, blue-flowered species possess a duplicated CYP75B lineage gene (CYP75B-D) with apparent 5′-hydroxylase activity, a derived D-type dihydroflavonol reductase (DFR), and a distinct anthocyanin biosynthetic pathway co-expression pattern. Phylogenetic analyses suggest that the D-type DFR pre-dated blue flower evolution, representing a permissive evolutionary shift later coupled with the evolution of CYP75B-D. Notably, blue flower evolution coincided with repeated pollinator transitions and major palaeoenvironmental change during the Middle Miocene. Together, these findings reveal an unexpected route to blue floral pigmentation and suggest that its evolution arose through the interaction of molecular innovation, regulatory evolution, and ecological opportunity.

## Introduction

Pollinator-mediated selection imposed by the differential preferences of pollinators favouring particular floral characteristics is a powerful force driving the evolution of floral traits (van der Niet and Johnson 2012; Schiestl and Johnson 2013; van der Niet et al. 2014). One such trait is flower colour, which plays a central role in attracting pollinators across the angiosperms by advertising potential rewards such as pollen or nectar (van der Niet and Johnson 2012; Lunau and Wester 2017). Specific colours often attract different pollinators, and shifts in floral colour driven by these preferences can lead to plant species divergence (Armbruster 2014).

Floral colour is primarily governed by specialised metabolites from three pigment classes: anthocyanins, betalains, and carotenoids (Grotewold 2006; Tanaka et al. 2008; Davies et al. 2012; Borghi and Fernie 2017; Wong et al. 2023). Anthocyanins are of key importance, with at least 600 compounds identified across three main structural forms: pelargonidin, cyanidin, and delphinidin. These compounds confer flowers the extraordinary ability to exhibit nearly the entire visible colour spectrum, as well as rare black hues (Iwashina 2015; Saigo et al. 2020; Nadot and Carrive 2021) and other dark tissue-specific floral structures (Wong et al. 2022a; Wong et al. 2022b; Kellenberger et al. 2023).

The anthocyanin biosynthesis pathway (ABP) is highly conserved and represents one of the best understood biochemical pathways in plants. Similarly, our understanding of the transcriptional regulators orchestrating diverse developmental stage-, tissue-, and stress-specific anthocyanin pigmentation programs in plants is also well-known (Albert et al. 2014). However, the recent discovery of a completely overlooked, yet critical dehydration step in anthocyanin biosynthesis (Eichenberger et al. 2023), highlights that there is still much to learn. Moreover, the insights gained thus far are based on a very limited fraction of flowering plant diversity (Wessinger and Rausher 2012; Larter et al. 2018; Larter et al. 2019; Wheeler et al. 2023; Marin-Recinos and Pucker 2024). Thus, it is likely that there are many unknown biosynthetic mechanisms yet to be discovered.

Loss of pigmentation often results in white and yellow flowers that account for the colour of approximately 44% of all flowers (Kattge et al. 2020; Dyer et al. 2021). Generally, inactivation mutations at one or more steps of the ABP often explain the loss of floral anthocyanins (Wessinger and Rausher 2012). Conversely, the chemical and genetic basis of natural floral hue shifts between red, orange, pink, and maroon to blue (7%) and purple (8%) flowers (and *vice versa*) remains a challenge to accurately predict based on flower colour alone.

Flavonoid 3’-hydroxylase (F3’H) and/or flavonoid 3’5’-hydroxylase (F3’5’H) are two crucial enzymes belonging to the plant cytochrome P450 75 (CYP75) multigene family that determine the flux of precursors across the three different branches of the anthocyanin pathway i.e. between red or orange-conferring pelargonidins, pink or maroon-cyanidins, and blue or purple-delphinidins (Tanaka et al. 2008; Davies et al. 2012; Seitz et al. 2015). Downstream of F3′H and F3′5′H, dihydroflavonols (dihydrokaempferol, DHK; dihydroquercetin, DHQ; or dihydromyricetin, DHM) are converted to their corresponding leucoanthocyanidins, flavan-3,3,4-triol, and anthocyanidins, via the action of dihydroflavonol reductase (DFR), leucoanthocyanidin dioxygenase/anthocyanidin synthase (ANS), and anthocyanin-related glutathione transferase (arGST) activities, respectively. Interestingly, DFR genes are known to evolve strict substrate specificity in certain plant lineages that have gained or lost the ability to produce delphinidin-rich flowers (Johnson et al. 2001; Smith et al. 2013; Larter et al. 2018; Wheeler et al. 2023; Sánchez-Cabrera et al. 2024; Liu et al. 2025).

There are three key mechanisms capable of inducing floral hue shifts: functional mutations affecting enzymes downstream of F3’H/F3’5’H; regulatory mutations that affect their expression (e.g. promoter changes); and functional mutations affecting specific transcriptional regulators (e.g. R2R3-MYB, bHLH) (Wessinger and Rausher 2012). Indeed, these three mechanisms underpin the few documented cases of interspecific floral colour transitions. These include shifts from delphinidin to pelargonidin-producing species in *Iochroma* (Solanaceae; Larter et al. 2018), and cyanidin-rich flowers of *Penstemon* (Plantaginaceae; Stone and Wessinger 2024) and *Aquilegia* spp. (Ranunculaceae; Edwards et al. 2021). Intraspecific examples include *Phlox drummondii* (Polemoniaceae; Hopkins and Rausher 2011) and *Lysimachia arvensis* (Primulaceae; Sánchez-Cabrera et al. 2021). In some cases, such as within the genera *Iochroma* and *Lysimachia*, multiple genetic changes in F3’5’H and DFR also underpin floral colour shifts (Smith et al. 2013; Larter et al. 2018; Sánchez-Cabrera et al. 2024).

The Orchidaceae is one of the largest and most diverse angiosperm families (Chase et al. 2015), with some estimates of up to ∼30,000 species. Rivalled only in size by the Asteraceae, the family is renowned for its staggering array of morphological forms, specialised pollination systems, and a reliance on mycorrhizal fungi for germination (Jersáková et al. 2006; Rasmussen and Rasmussen 2009; Schiestl and Schlüter 2009; Yukawa et al. 2009; Givnish et al. 2015; Bohman et al. 2016; Wong et al. 2017; Ackerman et al. 2023; Perkins et al. 2023; Freudenstein 2024). These characteristics make orchids an ideal group for addressing evolutionary questions, particularly with respect to the role of biotic interactions in speciation.

With more than 900 species, the predominantly Australian terrestrial orchid tribe Diurideae (Orchidoideae) is well-known for its floral colour diversity (Kores et al. 2001; Weston et al. 2014; Peakall et al. 2021; O’Donnell et al. 2024). While blue-, purple-, and violet-flowered species are rare across orchids, yet these colours are repeatedly observed across two subtribes within the tribe: the predominantly blue/violet-flowered Thelymitrinae encompassing the genera *Epiblema, Calochilus,* and *Thelymitra* (Bernhardt and Edens-Meier 2014; Nargar et al. 2018; Nauheimer et al. 2018); and the Caladeniinae, the largest subtribe within the Diurideae (Clements et al. 2015; Hopper and Brown, 2004).

Within the Caladeniinae alone, several distinct floral colour shifts are apparent across the phylogeny (**Figure 1**). The first is the emergence of a blue-flowered clade, amongst white- and maroon-flowered, food-rewarding relatives. This blue clade contains approximately 13 species split across the genera *Cyanicula*, *Elythranthera,* and *Glossodia* (Clements et al. 2015). Delphinidin-based anthocyanins are confirmed to underpin the floral colouration in *Elythranthera* and *Glossodia* (George et al. 1973; Wong et al. 2024), and are predicted to do so in *Cyanicula*. A subsequent shift from blue-coloured flowers to brightly coloured white, pink, and yellow species is observed in three *Caladenia* subgenera (*C.* subg. *Stegostyla*, *C.* subg. *Caladenia,* and *C.* subg. *Elevatae*) (Phillips et al. 2009, Xu et al. 2017, Zhou et al. 2025, Zhao et al. 2026). Finally, a shift towards predominantly maroon-flowered species occurs in the clade including *C.* subg. *Calonema* and *C.* subg. *Phlebochilus* which contains 250+ spp. (Hopper and Brown 2004; Clements et al. 2015; Peakall et al. 2021).

**Figure 1.**
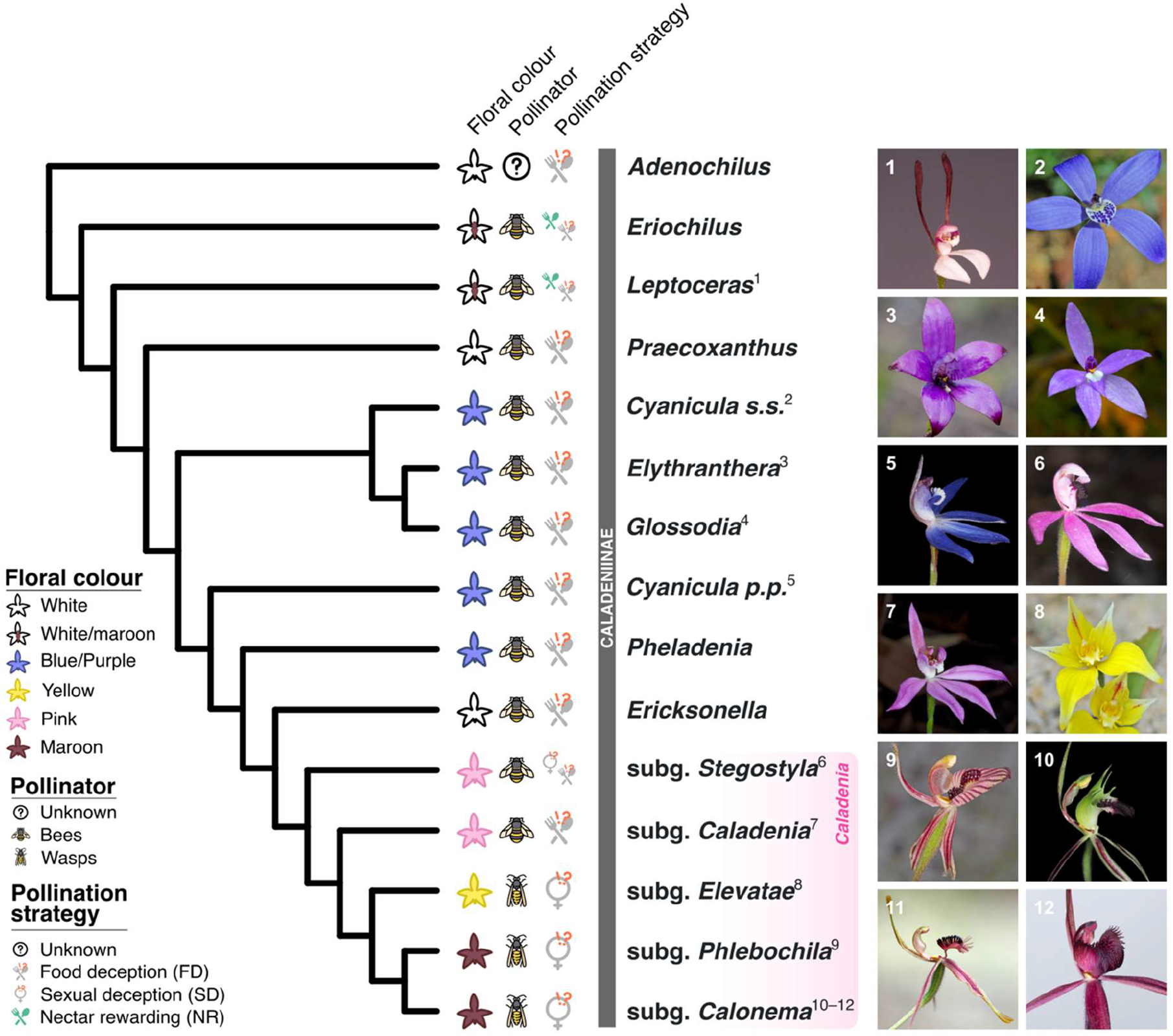
Summary topology of the Caladeniinae adapted from Peakall et al. (2021) and O’Donnell et al. (2024), with floral colour, pollinator, and pollination syndrome mapped as traits. Representative species photographs: 1. *Leptoceras menziesii, 2. Cyanicula gemmata* 3. *Elythranthera brunonis*, 4. *Glossodia major*, 5. *Cy. caerulea*, 6. *Caladenia congesta*, 7. *C. hillmanii*, 8. *C. flava*, 9. *C. cairnsiana*, 10. *C. atrovespa*, 11. *Caladenia barbarossa*, 12. *Caladenia concolor*. Photo credits: 1, 2, 3, 6, 8, 9, 11, 12 - R. Peakall; 5, 7, 10 - R.P. O’Donnell.

As an ideal study system, here we combined phylotranscriptomics, targeted metabolite profiling, and gene expression analysis to investigate the genetic and chemical basis of floral colour shifts across the Caladeniinae. We identified an unexpected route to blue pigmentation involving a duplicated CYP75B lineage, a derived D-type DFR and coordinated reorganisation of anthocyanin pathway expression. To contextualise these findings, we placed the observed floral colour shifts within a temporal and evolutionary framework, revealing a complex interplay between the micro- and macroevolutionary processes underpinning the evolution of blue flowers and their reversal.

## Methods

### Study system and sampling

The Caladeniinae comprises the genera *Adenochilus*, *Eriochilus*, *Leptoceras*, *Praecoxanthus*, and the species-rich genus *Caladenia*, which exhibits diverse floral colours and pollination strategies. We follow the taxonomy adopted by the Australian Plant Name Index (APNI) and Australian Plant Census (APC), which recognise *Cyanicula*, *Elythranthera*, and *Glossodia* as genera distinct from *Caladenia* (CANBR 2025). We sampled eight blue-flowered species from *Cyanicula*, *Elythranthera*, and *Glossodia*, together with 39 maroon-flowered *Caladenia* species and four outgroup taxa from the Diurideae and Cranichideae, representing a total of 51 species. Floral tissues were collected from naturally occurring populations during periods of pollinator activity, immediately snap-frozen in liquid nitrogen, and stored at −80°C until further use. Detailed information on taxon selection, pollination syndromes, sampling localities, voucher specimens, and tissue types is provided in the **Supplementary Methods and Supplementary Table S1**.

### Metabolite analysis

Anthocyanin pigment analysis followed the UHPLC-DAD-MS/MS methods of Wong et al. (2024). Briefly, floral tissues were homogenised and extracted in acidified methanol (70% methanol, 29% water, 1% formic acid) at 4°C for 24 h before centrifugation and analysis. Samples were analysed using a Thermo Scientific UltiMate 3000 UHPLC system coupled to a Q Exactive Plus Orbitrap mass spectrometer. Anthocyanins were identified using UV-visible absorbance spectra, exact mass measurements, MS/MS fragmentation patterns, relative elution order, and comparison with the KNApSAcK database (Afendi et al. 2012) and published literature. Relative anthocyanin abundance was estimated from integrated 520 nm peak areas using cyanidin 3-O-glucoside chloride as an external standard. Detailed chromatographic conditions, instrument settings, and compound identification procedures are provided in the Supplementary Methods. Targeted metabolite profiling using UHPLC–DAD–MS/MS was performed on several representative species of each major clade to establish the anthocyanin chemistry underpinning their floral colouration. This included two out of five outgroup species (*R. slateri, L. menziesii*), five out of the eight blue Caladeniinae species (*E. brunonis, E. emarginata, G. major, Cy. gertrudiae, Cy. caerulea*), *C. ustulata* (*C.* subg. *Stegostyla*, two *C.* subg. *Caladenia* species (*C. fuscata* and *C. carnea*), *C. nana* (*C.* subg. *Elevatae*), and 19 maroon *Caladenia* species (six *C.* subg. *Calonema* and 14 *C.* subg. *Phlebochilus*).

### Transcriptome sequencing

Preparation of samples for sequencing was performed following methods outlined by Wong et al. (2017; 2019), involving total RNA extraction with the Qiagen® RNeasy® Plant Mini Kit (Qiagen, Australia) with no deviations from the manufacturer’s protocol. To account for genetic variability and provide representative samples for each species, a minimum of five individuals were pooled to generate each species sample. Subsequent RNA quantitation and quality checks were conducted on an Agilent 2100 Electrophoresis Bioanalyzer (Agilent) using an RNA 6000 Nano chip. Poly(A) mRNA isolation and cDNA library construction was performed using NEBNext Poly(A) mRNA Magnetic Isolation Module and Ultra RNA Library Prep Kit for Illumina (NEB, Australia), respectively, as per the manufacturer’s instructions. PCR enrichment was run for 13 cycles. Paired-end (2 x 250bp) sequencing was performed using the Illumina NextSeq 500 platform (Illumina Technologies, USA) at the Biomolecular Resource Facility at the Australian National University (ANU) campus.

### *De novo* transcriptome assembly and functional annotation

Raw paired-end reads were quality filtered and adapter trimmed using fastp v0.23.0 (Chen et al. 2018) before *de novo* transcriptome assembly with Trinity v2.14.0 (Grabherr et al. 2011; Haas et al. 2013). Transcriptome completeness was assessed using BUSCO v5 (Waterhouse et al. 2018) with the embryophyta_odb10 database (**Supplementary Figure S1**). Redundant transcripts were filtered using the tr2aacds4 pipeline in EvidentialGene (Gilbert 2013) to generate species-specific reference transcriptomes. Functional annotation was performed using Mercator4/MapMan4 (Schwacke et al. 2019) and DIAMOND searches against UniProtKB (Buchfink et al. 2015). Candidate anthocyanin biosynthetic pathway (ABP) genes, including *CHS*, *CHI*, *F3H*, *F3’H*, *F3’5’H*, *FLS*, *DFR*, *ANS*, and *arGST*, were prioritised for downstream analyses (**Supplementary Figure S2**). Detailed software settings and filtering parameters are provided in the **Supplementary Methods**.

### Species and gene tree inference

Putative near-single-copy orthologs were identified across the 51 species using OrthoFinder v3.0 (Emms and Kelly 2019) with DIAMOND searches and MSA-based gene tree inference. Maximum-likelihood gene trees were subsequently inferred with IQ-TREE 2 v2.0.6 (Minh, Schmidt, et al. 2020) and used to construct a coalescent-informed species tree with ASTRAL III v5.5.4 (Zhang et al. 2018). Branch support was assessed using local posterior probabilities (Sayyari and Mirarab 2016), while gene and site concordance factors were estimated using IQ-TREE 2 (Minh, Hahn, et al. 2020). Trees were visualised and rooted in iTOL v6 (Letunic and Bork 2024). The timetree adapted from O’Donnell et al. (2024) was visualised using the R packages ggtree v3.13.0 (Yu et al. 2017) and deeptime v2.1.0 (Gearty 2024). Candidate CYP75 and DFR coding and protein sequences were aligned using MAFFT v7 with the L-INS-i algorithm (Katoh et al. 2002; Katoh and Standley 2013), and maximum-likelihood gene trees were inferred using IQ-TREE 2. Detailed software settings and command-line parameters are provided in the Supplementary Methods.

### Positive selection analyses

Signatures of positive selection in CYP75 and DFR were assessed using EasyCodeML v1.4 (Gao et al. 2019), a graphical interface for the CodeML program implemented in the PAML package (Yang 2007). Analyses were performed on reduced datasets comprising focal lineages and their closest orthologous/paralogous relatives rather than complete gene family phylogenies, thereby targeting specific evolutionary transitions of interest while minimising the effects of excessive sequence divergence. Detailed taxon sampling, foreground branch designations, and model settings are provided in the **Supplementary Methods**.

### Anthocyanin biosynthetic pathway gene expression analysis

Paired-end reads were mapped to the filtered reference transcriptomes using Bowtie2 v2.4.5 (Langmead and Salzberg 2012), and read counts were summarised with FeatureCounts v2.0.4 (Liao et al. 2014). Using the tissue type most representative of floral colour for each species, fragments per kilobase of transcript per million mapped reads (FPKM) were calculated for annotated anthocyanin biosynthetic pathway (ABP) genes and log₂-transformed prior to analysis. Expression differences between blue- and maroon-flowered species were assessed using non-parametric Wilcoxon rank-sum tests (Mann-Whitney U tests) implemented in the R package *stats* v4.4.1 (R Core Team 2024). Detailed mapping, filtering, and summarisation parameters are provided in the **Supplementary Methods**.

### Phylogenetic principal component analysis

Phylogenetic principal component analysis (pPCA) was used to investigate relationships between anthocyanin biosynthetic pathway (ABP) gene expression and floral anthocyanin chemistry while accounting for shared evolutionary history among species. Analyses were performed using the phyl.pca function in *phytools* v2.3.0 (Revell 2012), and phylogenetic correlation matrices were extracted to examine gene expression relationships among cyanidin- and delphinidin-producing taxa. To assess both strong and weak phylogenetic structure in the data, phylogenetically aligned component analysis (PACA) was performed using the gm.prcomp function in *geomorph* v4.0.8 (Adams and Otárola-Castillo 2013; Collyer and Adams 2021).

## Results

### Phylotranscriptome of the Caladeniinae

The ASTRAL summary species tree for 51 Caladeniinae species was inferred using 939 ML gene trees, with *Pterostylis curta* (Cranichideae) used to root the tree. *Leptoceras menziesii* was inferred as sister to all other Caladeniinae species included in this study (**Figure 2**). Within the clade containing all other Caladeniinae species were two main clades; the first, a well-supported clade on a long branch (LPP = 1.0; gCF = 45; sCF = 60) comprised of the blue-flowered genera *Cyanicula, Elythranthera,* and *Glossodia*; and the second, a clade containing an additional blue-flowered clade encompassing the eastern Australian *Cyanicula* species, *Cy. caerulea* and the Western Australian *Cy. aperta*, sister to a large clade containing all maroon-coloured *Caladenia* species corresponding with *C.* subg. *Stegostyla* (*C. ustulata* and *C. congesta*), *C.* subg. *Caladenia* (*C. fuscata* and *C. carnea*), *C.* subg. *Elevatae* (*C. nana* and *C. flava*), *C.* subg. *Calonema* (9 species), and *C.* subg. *Phlebochilus* (24 species). This topology is broadly congruent with previous studies (**Figure 2A – 2C**) (Kores et al. 2001; Clements et al. 2015; Wong and Peakall 2022; O’Donnell et al. 2024). As in previous phylogenomic studies (Peakall et al. 2021), species-level relationships within the *Calonema* and *Phlebochilus* clades were characterised by low LPP values and low gCF/sCF scores (LPP = 0.00–1.0, 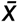 = 0.65; gCF = 0.00– 48, 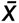 = 11; sCF = 30 – 82, 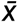 = 44).

**Figure 2.**
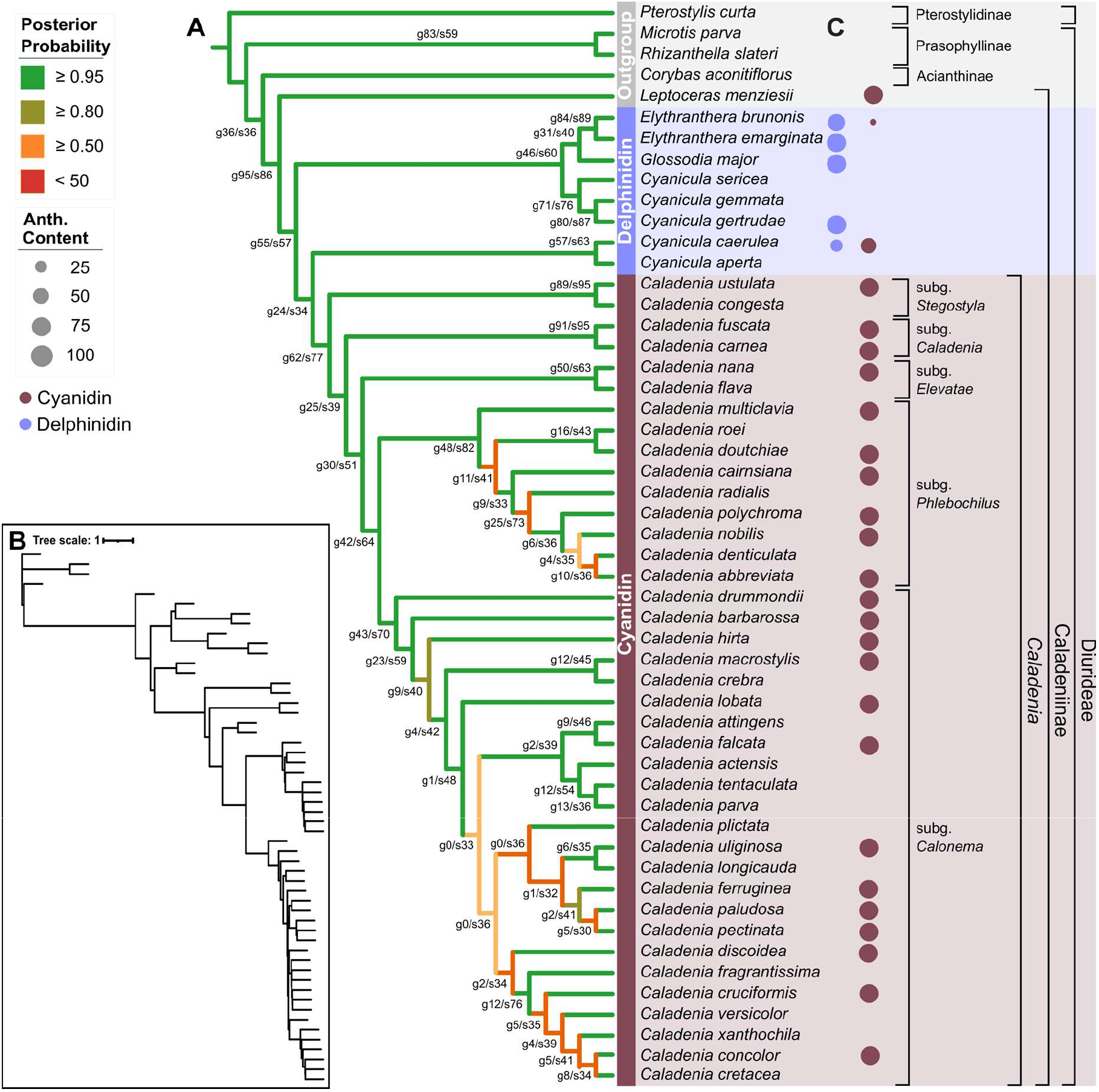
Phylotranscriptomic framework and floral anthocyanin chemistry of the Caladeniinae. **(A)** ASTRAL species tree inferred from 939 orthogroups spanning 51 species of Diurideae and the outgroup Pterostylis curta. Branch colours indicate local posterior probability (LPP) support, while node labels show LPP/gCF/sCF values. **(B)** Phylogram of the ASTRAL species tree with branch lengths expressed in coalescent units. **(C)** Distribution of anthocyanin chemistry across representative taxa. Circle colour indicates the predominant anthocyanin class, with purple representing delphinidin-derived pigments and maroon representing cyanidin-derived pigments. Circle size is proportional to the relative abundance of each anthocyanin class.

### Floral anthocyanin chemistry of the Caladeniinae

We found that the white petals with pink to dark red markings of *Leptoceras menziesii* flowers accumulate only two cyanidin-based pigments, cyanidin glucoside and cyanidin dimalonyl glucoside (**Figure 3**). Flowers of almost all blue Caladeniinae species were underpinned by a more complex delphinidin-based anthocyanin pigment profile. *Elythranthera brunonis* and *Cy. caerulea* were two exceptions, with flowers accumulating both delphinidin- and cyanidin-based pigments at a 9:1 and 3:2 ratio, respectively (**Figure 2C**). Conversely, all 38 species with maroon-coloured flowers exclusively produced cyanidin-based anthocyanins. These patterns indicate that the gain of blue floral pigmentation in Caladeniinae is consistently associated with the evolution of delphinidin biosynthesis (**Figure 3, Supplementary Figure 3 and 4**).

**Figure 3.**
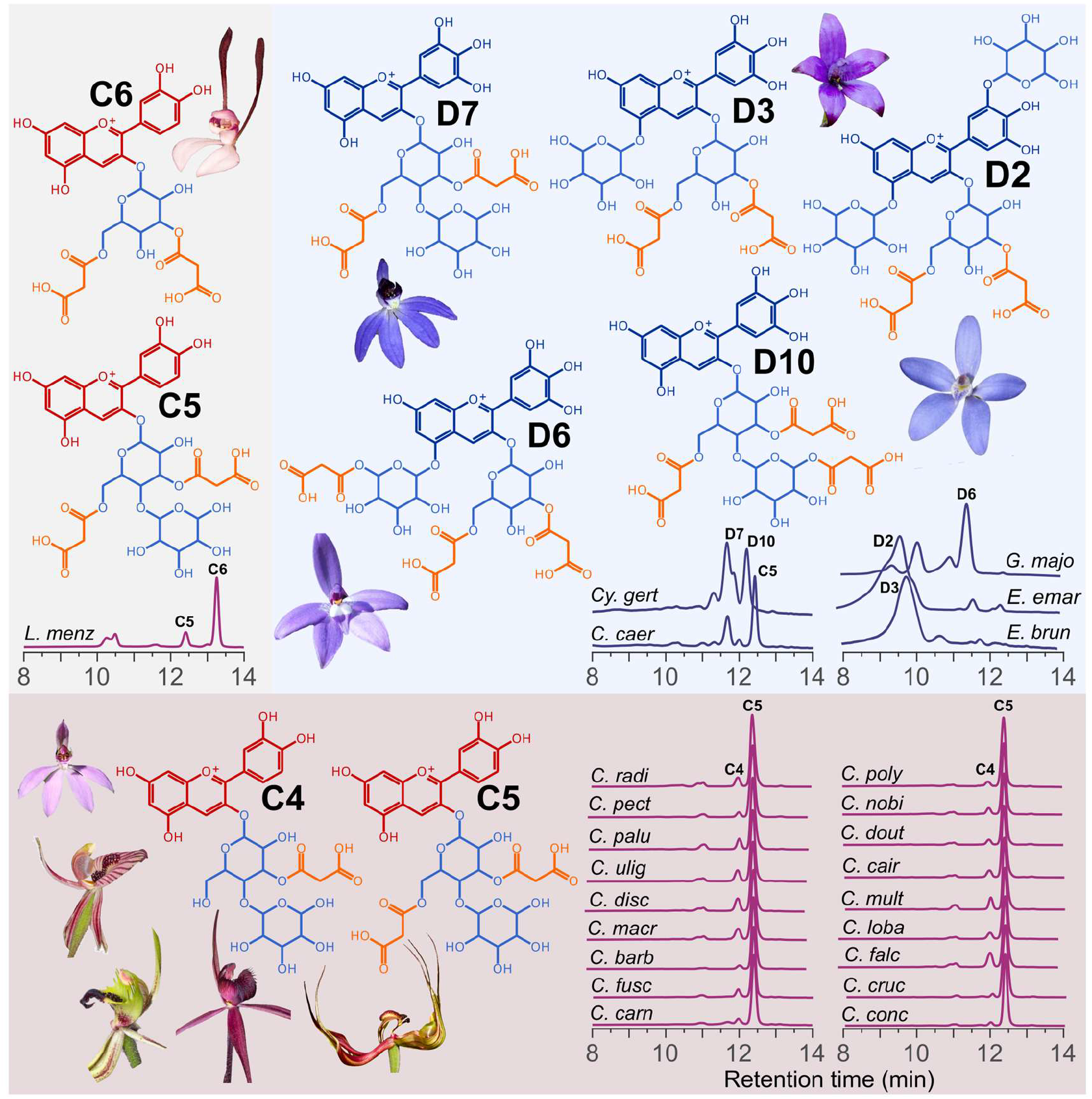
Anthocyanin profiles and putative structures across representative Caladeniinae taxa. Taxa were selected to broadly represent the phylogenetic diversity outlined in Figure 2. For each representative taxon, the chromatogram is shown together with the putative anthocyanin structure corresponding to the major peak. Putative structural assignments were based on precursor ion masses and MS/MS fragmentation patterns. See **Supplementary Figures S3 and S4** for further information.

**Figure 4.**
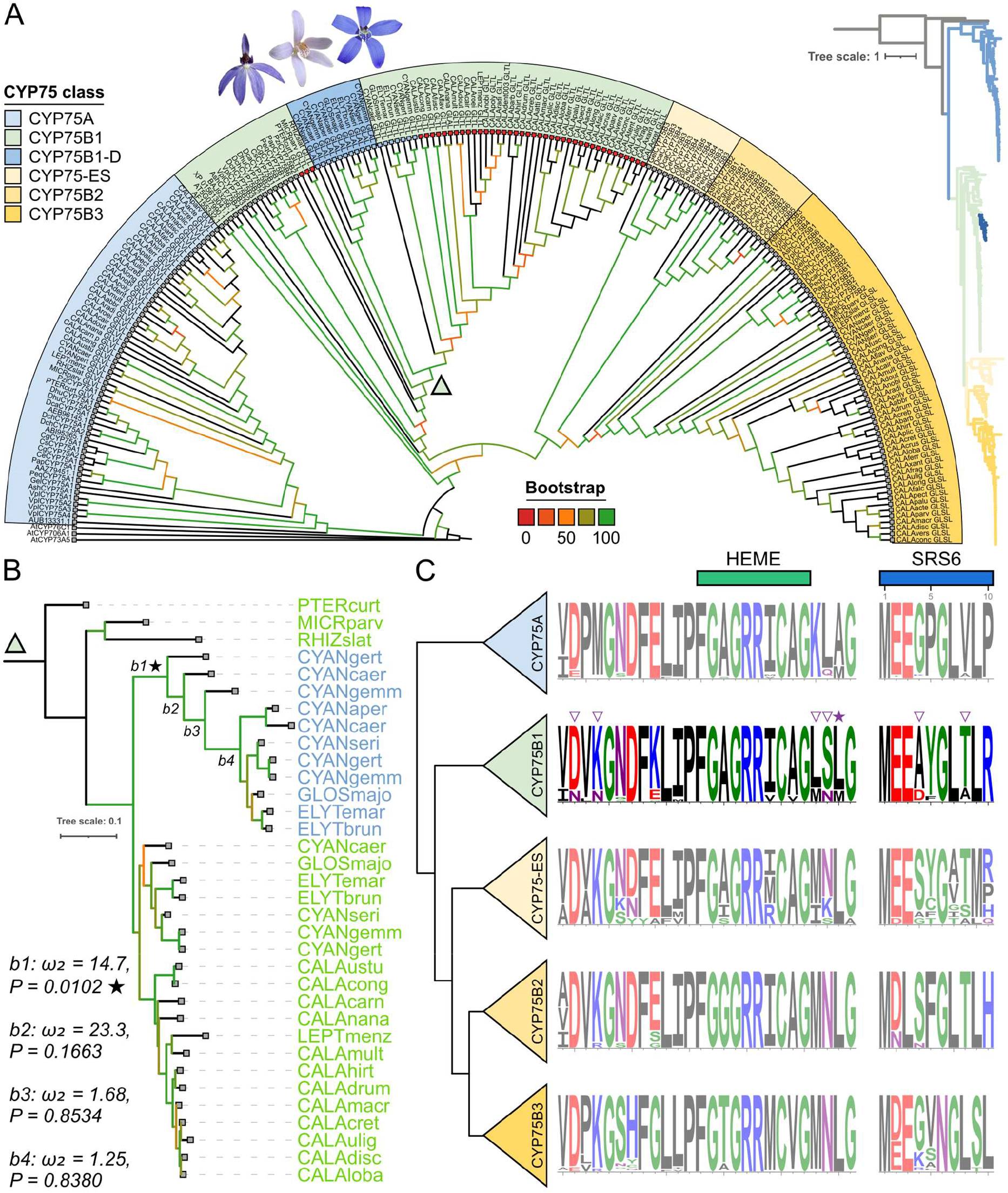
Evolutionary history of CYP75 genes and signatures of selection associated with the origin of the CYP75B-D lineage. **(A)** Maximum-likelihood phylogeny of CYP75 proteins identified from published orchid genomes and study species. The outgroup comprises representative CYP sequences from non-CYP75 families. Branch colours indicate bootstrap support values. CYP75B-ES denotes an Epidendroideae-specific CYP75B lineage, whereas CYP75B-D denotes the duplicated CYP75B lineage associated with delphinidin-producing Caladeniinae taxa. Flower photographs (left to right: *Cy. caerulea*, *Cy. gertrudiae*, and *Cy. gemmata*) illustrate representative species possessing two CYP75B-D paralogs. Square symbols at the tips of the CYP75B1 clade indicate floral anthocyanin chemistry, with purple and maroon denoting delphinidin- and cyanidin-producing taxa, respectively. An inset phylogram (right) displays branch lengths, and the complete gene tree is provided in **Supplementary Figure S5**. **(B)** Reduced CYP75B phylogeny used for positive selection analyses. Foreground branches tested using EasyCodeML are indicated as b1 – b4. Summary of branch-site analyses results are shown, with D1 representing the only branch exhibiting a significant signal of positive selection. **(C)** Sequence logos showing amino acid conservation across major CYP75 clades for the substrate recognition site 6 (SRS6) region and the conserved cytochrome P450 haem-binding motif. Filled stars indicate residues with BEB posterior probabilities >0.95, whereas open triangles indicate residues with BEB posterior probabilities of 0.93 – 0.95 from branch-site analyses. Residue height is proportional to amino acid conservation at each position.

### Phylogenetic analysis reveals a divergent CYP75B lineage associated with blue pigmentation in Caladeniinae

To gain insights into the genetic basis underpinning floral colour shifts from maroon to blue and the reversal back to maroon-flowered species, we first investigated the evolutionary history of the CYP75 gene family within the Caladeniinae. Given the established role of *CYP75A* genes in 3′,5′-hydroxylation and blue flower pigmentation across angiosperms, we initially expected blue-flowered species to possess a lineage-specific CYP75A orthologue. To test this hypothesis, we performed a comprehensive genome-wide analysis of the CYP75A and CYP75B gene families across all 13 published orchid genomes and the Caladeniinae, building on the phylogenetic framework of Li et al. (2023). From the transcriptome assemblies generated in this study, we recovered more than 120 CYP75 sequences comprising 32 *CYP75A* (candidates) and 97 *CYP75B* candidates (**Figure 4, Supplementary data S1**).

Phylogenetic analysis revealed that 32 sampled Australian species expressed a candidate CYP75A. Representative CYP75A genes were missing for outgroup species *Corybas aconitiflorus,* six of the eight blue Caladeniinae species, and 13 maroon *Caladenia* species. However, phylogenetic placement of these CYP75A genes corresponded closely to the species relationships of the broader Orchidaceae phylogeny and aligned with the species tree produced in this study (**Figure 2**). The remaining 97 sequences were placed confidently into the CYP75B subgroup across three distinct clades (**Figure 4A**), with phylogenetic relationships largely consistent with that of the species tree. The first orchid CYP75B clade (arbitrarily labelled CYP75B1) included a representative for all study species, excluding *Cy. aperta*, which was absent from its respective *de novo* transcriptome assembly.

Unexpectedly, a small, highly divergent CYP75B lineage (hereafter CYP75B-D) was recovered within the broader CYP75B1 clade and included representatives from all eight blue-flowered Caladeniinae species (**Figure 4A**). This lineage was separated from canonical CYP75B1 sequences by a long branch, indicative of substantial amino-acid divergence consistent with an ancestral gene duplication followed by sequence diversification. Multiple CYP75B-D paralogs were recovered from the transcriptome assemblies of *Cy. gertrudae*, *Cy. caerulea* and *Cy. gemmata*, whereas the remaining blue-flowered species possessed a single CYP75B-D copy. Notably, corresponding CYP75B1 copies were also retained in all blue-flowered Caladeniinae species, indicating that the divergent CYP75B-D lineage evolved alongside the ancestral CYP75B1 paralog rather than replacing it. The strong association of CYP75B-D with blue-flowered taxa contrasted sharply with the broader CYP75A and CYP75B phylogenetic patterns. No Caladeniinae sequences matching the epidendroid-specific CYP75B-ES or orchid CYP75B2 clades were detected. In contrast, the orchid CYP75B3 clade included representatives from 40 of the 51 study species. Sequences were absent for the outgroup species *Corybas aconitiflorus* and *Pterostylis curta*, as well as four of the eight blue-flowered Caladeniinae species (*E. brunonis*, *E. emarginata*, *G. major* and *Cy. gemmata*). Representation was also absent for six of the 38 maroon-flowered *Caladenia* species.

### Acquisition of a CYP75A-like SRS6 motif within a duplicated CYP75B lineage

Previous studies have identified position 8 within substrate recognition site 6 (SRS6) as a key determinant of catalytic specificity in CYP75 enzymes, distinguishing flavonoid 3′-hydroxylase (F3′H; CYP75B) from flavonoid 3′,5′-hydroxylase (F3′5′H; CYP75A) activity (Seitz et al. 2007; Seitz et al. 2015). Alanine (Ala) at this position is strongly associated with 5′-hydroxylation activity characteristic of CYP75A enzymes, whereas threonine (Thr) is typically observed in CYP75B enzymes exhibiting only 3′-hydroxylation activity. More recent studies have demonstrated that CYP75B enzymes can acquire F3′5′H activity through relatively simple amino acid substitutions, including replacement of Thr with Ala at position 8 of SRS6 (Seitz et al. 2006; Seitz et al. 2007).

To investigate whether similar functional shifts may have occurred within the Caladeniinae, we examined amino-acid variation across the SRS6 region of all recovered CYP75 sequences (**Figure 4B**). Consistent with Li et al. (2023), all candidate Caladeniinae CYP75A proteins possessed valine (Val) at position 8 of SRS6. Likewise, canonical Caladeniinae CYP75B1 proteins retained the conserved Thr residue characteristic of enzymes with F3′H activity (**Supplementary Data S2**). On the other hand, all members of the divergent CYP75B-D lineage possessed Ala at position 8 of SRS6 despite being phylogenetically nested within the CYP75B1 clade. This substitution is in the same residue previously demonstrated to confer 5′-hydroxylation capability in CYP75B enzymes (Seitz et al. 2006; Seitz et al. 2007; Lui et al. 2020). Consistent with this observation, all species possessing CYP75B-D accumulated delphinidin-derived anthocyanins (**Supplementary Figure 3**), supporting the hypothesis that CYP75B-D functions as a *bona fide* flavonoid 3′,5′-hydroxylase despite its CYP75B lineage. A notable exception was the maroon-flowered species *C. nana*, which lacked a CYP75B-D paralog but possessed Ala at the same SRS6 position within a canonical CYP75B1 copy. Despite carrying the derived Ala residue, *C. nana* accumulated only cyanidin-derived anthocyanins and no detectable delphinidin (**Supplementary Figure 4**), indicating that the substitution alone in the standard non-duplicated gene is insufficient to confer the blue-pigmentation phenotype. Additionally, members of the CYP75B3 clade possessed serine (Ser) at position 8, rather than Thr or Ala. In contrast to CYP75B-D, however, the distribution of this substitution was not strongly associated with delphinidin accumulation or blue floral pigmentation in the Caladeniinae. Collectively, these results revealed a strong association between delphinidin production and the presence of the derived CYP75B-D lineage carrying the Ala-containing SRS6 motif, whereas cyanidin-based pigmentation was largely associated with canonical CYP75B1 proteins retaining the ancestral Thr residue.

### Signatures of positive selection during the emergence of the CYP75B-D lineage

To test whether the emergence of the CYP75B-D lineage was likely accompanied by adaptive protein evolution, branch-site models were applied to a representative CYP75B1 phylogeny while minimising redundancy among closely related taxa. Significant evidence of episodic positive selection was detected only on the branch corresponding to the origin of the CYP75B-D lineage (b_1_; ω₂ = 14.7, *P* = 0.012; **Figure 4B**), whereas no evidence of positive selection was detected on subsequently derived branches (b_2_ – b_4_; *P* = 0.166-0.853; **Supplementary data S3**). Bayes Empirical Bayes (BEB) analyses identified two codons with strong support for positive selection, corresponding to residues 385A (PP = 0.990) and 436L (PP = 0.959) (**Figure 4B**). Several additional residues exhibited elevated posterior support (PP > 0.90), including 434L, 435S, 468A and 472T. Notably, residue 472 corresponds to position 8 within SRS6, the key site underpinning the transition from the ancestral GLTL motif to the derived GLAL motif associated with 5′-hydroxylation activity. Candidate adaptive residues were concentrated around the heme-binding region and SRS6, suggesting that adaptive sequence diversification accompanied the emergence of CYP75B-D and its associated delphinidin-producing phenotype in blue-flowered Caladeniinae (**Figure 4C**).

### Evolution of Asn (N)- and Asp (D)-type dihydroflavonol 4-reductase substrate-binding motifs in Caladeniinae

Dihydroflavonol 4-reductase (DFR) occupies a key branch point in anthocyanin biosynthesis, and evolutionary changes in its substrate-binding domain underpins altered anthocyanin composition and contribute to floral colour diversification (Johnson et al. 2001; Petit et al. 2007; Smith and Rausher 2011; Wheeler et al. 2022). Previous studies have shown that substrate specificity is strongly influenced by the third residue within a conserved 26-amino-acid substrate-binding region, resulting in N-type, D-type and non-N/non-D DFR variants (Johnson et al. 2001). Given the close association between blue floral pigmentation and delphinidin accumulation in the Caladeniinae, we investigated whether the gain of blue pigmentation was accompanied by the evolution of a distinct DFR substrate-binding sites.

DFR sequences were recovered for 48 of the 51 study species (**Supplementary Data S4**) and shared 87-97% amino acid identity with the functionally characterised *Glossodia major* DFR (mean 89.3%; Wong et al. 2024). Phylogenetic analysis revealed strong clustering of blue-flowered Caladeniinae and maroon-flowered *Caladenia* species, with sequences within each clade sharing 95.6% and 98.1% amino acid identity, respectively (**Figure 5A**). The only exceptions were the partial *DFR* sequence recovered from the yellow-flowered *C. flava* and the absence of detectable DFR transcripts from the pale-flowered species *C. longicauda* and *C. cretacea*, all of which exhibit reduced anthocyanin accumulation. Interestingly, despite lacking much of the C-terminal coding region, the partial *C. flava DFR* sequence was nested within the blue-flowered lineage.

**Figure 5.**
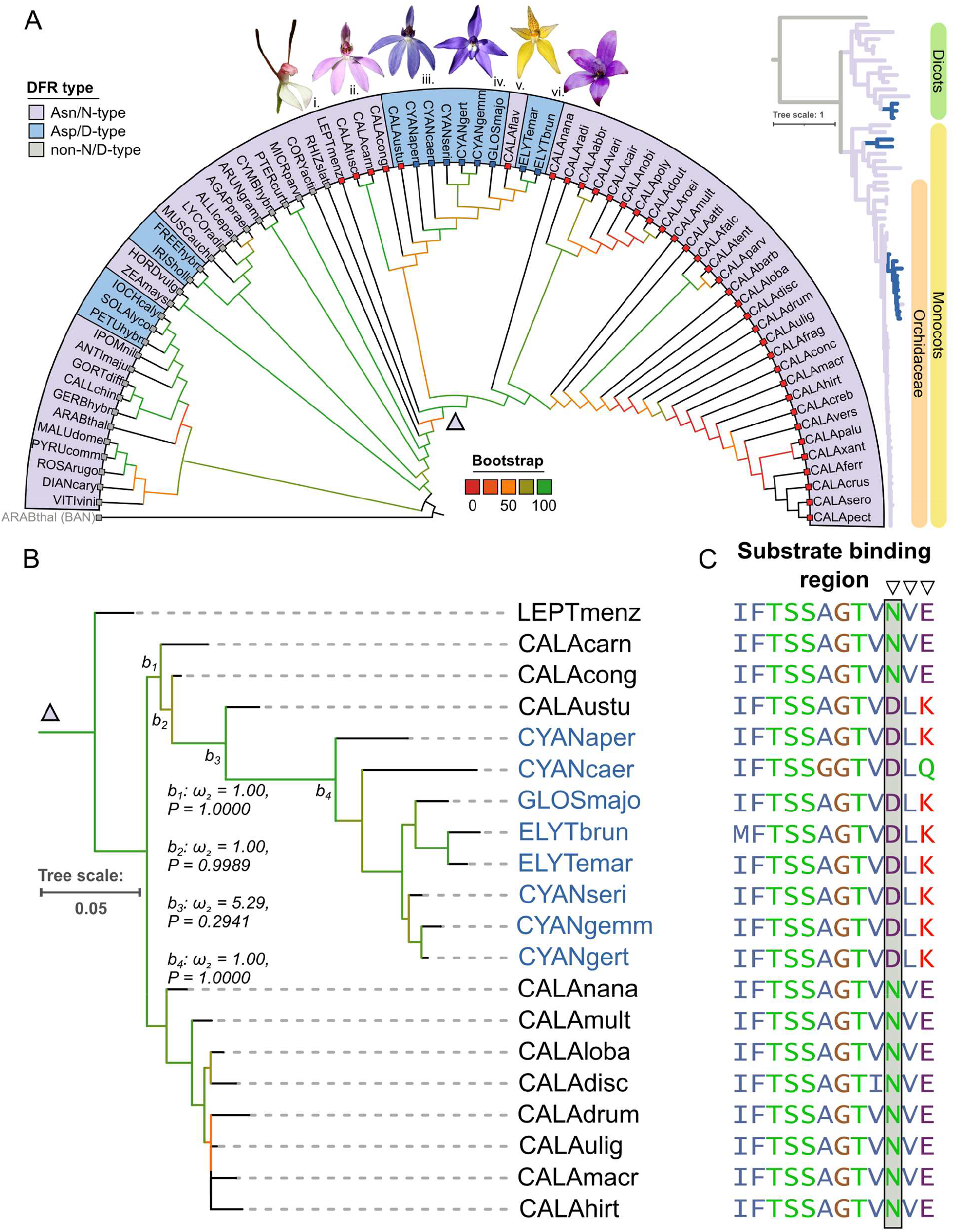
Evolutionary history of DFR and signatures of selection associated with the origin of D-type DFRs in the Caladeniinae. **(A)** Maximum-likelihood phylogeny of DFR amino acid sequences from all study species. Branch colours indicate substrate-binding residue classes at the diagnostic position, with D-type (Asp), N-type (Asn), and non-D/non-N variants shown according to the colour key. Branch support values are represented by bootstrap support. Flower photographs (left to right: *Leptoceras menziesii*, *Caladenia carnea*, *Cyanicula caerulea*, *Glossodia major*, *Caladenia flava*, and *Elythranthera brunonis*) illustrate representative taxa spanning the inferred transition from ancestral N-type to derived D-type DFRs within the small finger orchid lineage and highlight the occurrence of *C. flava* within the D-type lineage despite the absence of blue floral pigmentation. An inset phylogram (right) displays branch lengths, and the complete gene tree is provided in **Supplementary Figure S6**. **(B)** Reduced DFR phylogeny used for positive selection analyses. Foreground branches tested using EasyCodeML are indicated, together with the results of branch-site analyses. **(C)** Alignment of the DFR substrate-binding region highlighting amino acid variation across major DFR lineages. Open triangles indicate sites identified by branch-site analyses with Bayes Empirical Bayes (BEB) posterior probabilities of 0.85–0.95.

Inspection of the conserved substrate-binding region revealed a striking clade-specific amino acid substitution. Representative Diurideae outgroups and all maroon-flowered *Caladenia* species possessed the ancestral Asparagine residue (N-type), whereas all blue-flowered Caladeniinae species possessed Aspartic acid (D-type) at the same position (**Figure 5B**; **Supplementary Data S5**). This substitution resulted from a single nucleotide polymorphism converting the codon AAT to GAT. Interestingly, the D-type lineage was sister to a small N-type clade comprising *C. carnea*, *C. fuscata* and *C. congesta*, placing the N-to-D transition near the base of the blue-flowered lineage. Unlike Caladeniinae CYP75B-D lineages; however, D-type DFRs did not form a duplicated lineage and instead evolved within the canonical DFR phylogeny. A notable exception was the maroon-flowered *C. ustulata*, which was positioned outside the blue-flowered clade despite also possessing a D-type DFR. Consistent with its maroon floral phenotype, anthocyanin analyses of *C. ustulata* detected only cyanidin-derived anthocyanins (**Supplementary Figure 4**).

To test whether the emergence of the D-type DFR lineage was accompanied by adaptive protein evolution, branch-site models were applied to a representative *DFR* phylogeny spanning the N-type and D-type lineages (**Figure 5C**). In contrast to CYP75B-D, no branch tested showed significant evidence of positive selection (all *P* > 0.05; **Supplementary data S6**). Across alternative foreground branch definitions, branch-site models either converged on the null model (b1, b2, b4: ω₂ = 1.0; *P* = 0.999-1.000) or yielded non-significant evidence of positive selection (b3: ω₂ = 5.29; *P* = 0.294). Bayes Empirical Bayes analyses similarly identified no codons exceeding the conventional significance threshold (PP > 0.95), although the diagnostic N-type/D-type residue at codon 141 exhibited elevated posterior support (PP = 0.843).

### Evolution of anthocyanin pathway gene expression associated with blue floral pigmentation in Caladeniinae flowers

We hypothesise that the gain of blue floral pigmentation in the Caladeniinae was accompanied by substantial alterations across the anthocyanin pathway, including the emergence of CYP75B-D, the evolution of D-type DFRs, and accumulation of delphinidin-derived anthocyanins. We therefore investigated whether these alterations were also associated with coordinated changes in ABP gene expression. To characterise pathway-wide expression evolution, we mapped the transcriptomic morphospace of core ABP genes using Phylogenetic Principal Component Analysis (phyloPCA) (**Figure 6**). Species separated strongly along the first two axes, which together explained 65.1% of the total variance (PC1 = 52.3%; PC2 = 12.8%). Blue-flowered species occupied a distinct region of transcriptomic space that was consistently separated from both ancestral and derived maroon-flowered lineages (**Figure 6A**). This pattern was further supported by Phylogenetically Aligned Component Analysis (PACA), where the Ordinary Least Squares (OLS)-centred PACA aligned the primary axis of phylogenetic signal with floral colour, and the Generalized Least Squares (GLS)-centred PACA retained the separation between blue- and maroon-flowered phenotypes after accounting for shared ancestry (**Supplementary Figure S7**).

**Figure 6.**
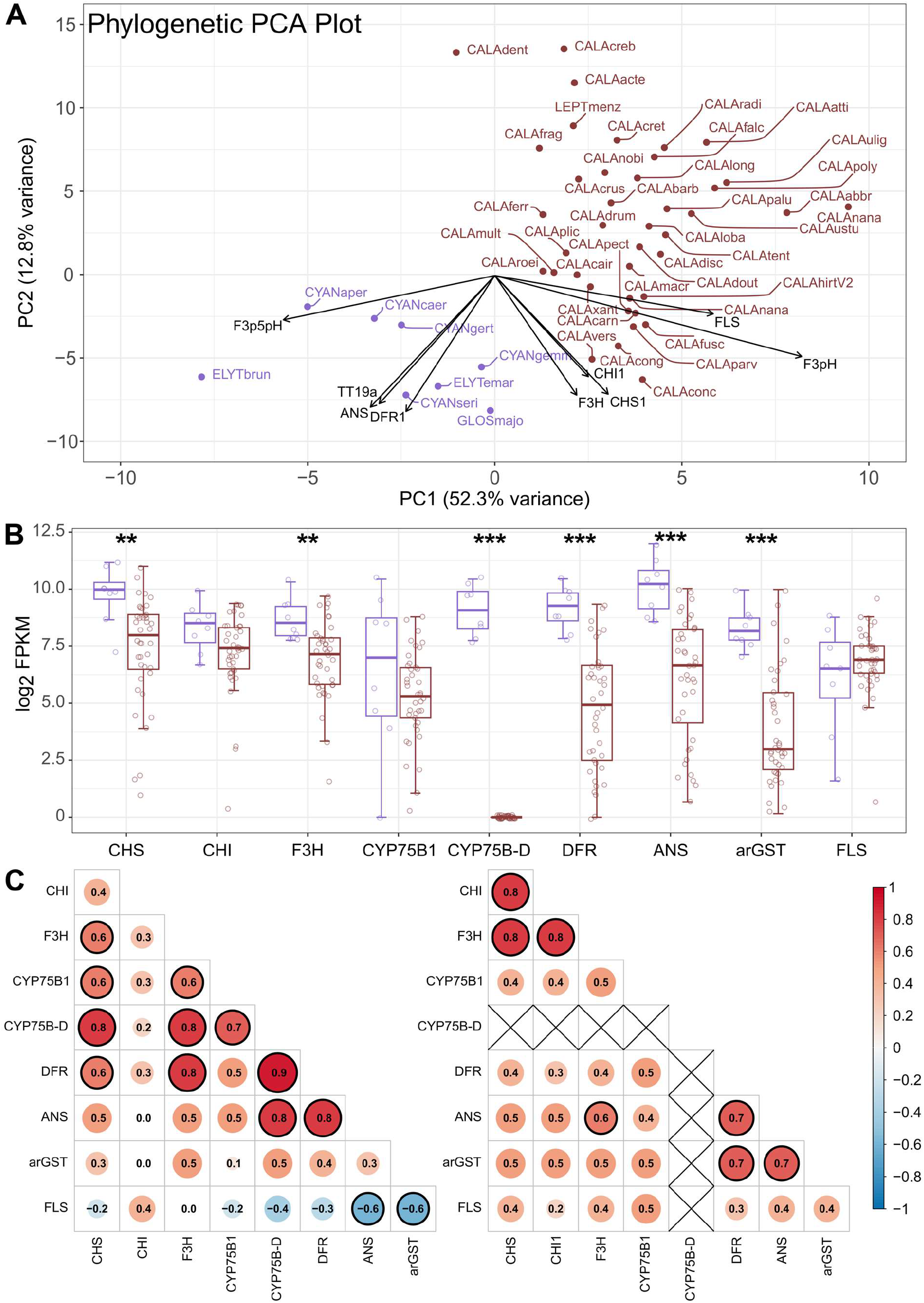
Evolution of anthocyanin biosynthetic pathway (ABP) expression and co-expression architecture in the Caladeniinae. **(A)** Phylogenetic principal component analysis (pPCA) of anthocyanin pathway gene expression based on log₂-transformed FPKM values. Species are coloured according to pigment class, with delphinidin-producing taxa shown in purple and cyanidin-producing taxa shown in maroon. Gene loadings on PC1 and PC2 indicate the contribution of each variable to the observed phylogenetic structure. **(B)** Comparison of ABP gene expression between cyanidin- and delphinidin-producing taxa. Group differences were assessed using non-parametric Wilcoxon rank-sum (Mann-Whitney U) tests. Asterisks indicate statistically significant differences (*P* < 0.05, **P** < 0.01, ***P*** < 0.001). **(C)** Pairwise gene expression correlation matrices for delphinidin- and cyanidin-producing taxa. Red, white, and blue indicate positive, neutral, and negative correlations, respectively. Circle size and colour intensity correspond to correlation strength, with Pearson correlation coefficients (*r*) shown within circles. Strong correlations (|*r*| ≥ 0.6) are highlighted by bold outlines. Abbreviations: **CHS**, chalcone synthase; **CHI**, chalcone isomerase; **F3H**, flavanone 3-hydroxylase; **FLS**, flavonol synthase; **CYP75B1**, flavonoid 3′-hydroxylase; **CYP75B-D**, duplicated CYP75B lineage; **DFR**, dihydroflavonol 4-reductase; **ANS**, anthocyanidin synthase; **arGST**, anthocyanin-related glutathione S-transferase.

The distinct positioning of blue- and maroon-flowered taxa within the transcriptomic morphospace was driven by polarised loadings of several key ABP genes (Figure 5A). *DFR*, *ANS*, *arGST* and *CYP75B-D* exhibited strong positive loadings associated with the blue-flowered lineage, whereas *CHS*, *CHI*, *F3H*, *CYP75B1* and *FLS* were associated with the maroon-flowered lineage, collectively distinguishing the transcriptomic signatures of delphinidin- and cyanidin-producing species. Notably, *CYP75B-D*, the duplicated lineage linked to delphinidin production, was among the strongest contributors to separation along PC1. Consistently, transcript abundance analyses revealed significantly higher expression of *CHS*, *F3H*, *FLS*, *CYP75B-D*, *DFR* and *ANS* in blue-flowered species relative to maroon-flowered lineages (**Figure 6B**). No significant differences were detected for *CHI*, *F3’H* or *arGST* expression between the two groups. Together, these patterns reveal a coordinated shift in both upstream and downstream ABP gene expression associated with the transition between cyanidin- and delphinidin-producing lineages.

These divergent expression profiles prompted a clade-specific co-expression analysis to determine whether the underlying architecture of the ABP also differed between blue- and maroon-flowered lineages (**Figure 6C**). Within the blue delphinidin-accumulating lineage, moderate-to-strong positive correlations (0.6 *r* < 0.8) connected *CHS*, *F3H*, *F3’H*, *CYP75B-D* and *DFR* into a highly integrated module. In contrast, *FLS* exhibited a strong negative correlation (*r* ≈ −0.6) with both *ANS* and *arGST*, suggesting a separation between competing flavonol- and anthocyanin-associated components of the pathway. The maroon cyanidin-producing lineage exhibited a markedly different co-expression architecture. Rather than a single integrated module, expression was partitioned into two distinct regulatory groups consisting of an early pathway module (*CHS*, *CHI* and *F3H*) and a late pathway module (*ANS*, *DFR* and *arGST*). Correlations among the remaining pathway components were comparatively weak (0.3 *r* < 0.5), indicating a less integrated network structure than that observed in the blue lineage. Collectively, these results reveal contrasting anthocyanin pathway architectures between blue and maroon Caladeniinae, with the blue lineage characterised by a more tightly coordinated expression network linking *CYP75B-D* to core anthocyanin biosynthetic genes.

## Discussion

### Phylotranscriptomics provides a robust new framework for understanding Caladeniinae evolution

Our phylotranscriptomic reconstructions recovered relationships broadly congruent with recent phylogenomic studies of the Diurideae, while substantially increasing species representation across the major Caladeniinae lineages (Peakall et al. 2021; O’Donnell et al. 2024). Support values were generally higher than those obtained from earlier studies based on a limited number of nuclear and plastid markers (Swarts et al. 2014; Weston et al. 2014; Clements et al. 2015), demonstrating the power of transcriptome-scale datasets for resolving relationships within this diverse orchid subtribe. Nevertheless, relationships within *C.* subg. *Calonema* and subg. *Phlebochilus* remained poorly resolved, and were characterised by short branches, low concordance among genes and sites, and reduced node support.

Previous studies have suggested that incomplete lineage sorting (ILS) associated with rapid diversification, hybridisation, or a combination of both may contribute to these patterns of discordance (Swarts et al. 2014; Peakall et al. 2021). Although O’Donnell et al. (2024) identified ILS as a major driver of discordance within the Diurideae backbone, its role within the Caladeniinae remains untested. Likewise, while hybridisation is frequently invoked in taxonomic treatments of the Caladeniinae (Hopper and Brown 2004; Clements et al. 2015), there is currently little molecular evidence supporting widespread natural hybridisation or extensive hybrid swarms within the group.

An alternative explanation may lie in the pollination biology of many members of subg. *Calonema* and *Phlebochilus* which predominantly engage in sexual deception (Peakall 2023). Members of these clades rely on highly specialised pollinator interactions (Peakall et al. 2021), where reproductive isolation can evolve through relatively small genetic changes affecting floral scent chemistry and pollinator attraction (Schiestl and Ayasse 2002; Xu et al. 2011; Peakall et al, 2010; Peakall and Whitehead 2014; Sedeek et al. 2014; Russo et al. 2024). Under this model, species boundaries may arise through changes at relatively few loci despite otherwise limited genomic divergence. The short branches and pervasive discordance observed here may therefore reflect genuinely shallow genomic differentiation among putative species rather than methodological limitations, highlighting the challenges of reconstructing evolutionary relationships in systems where reproductive isolation is driven by highly localised genetic changes.

### Gene duplication of CYP75B and recruitment of F3’5’H activity underpins delphinidin accumulation in blue-flowered Caladeniinae species

Floral colour evolution in the Caladeniinae is marked by transitions from ancestral maroon to derived blue flowers, followed by a subsequent reversion to maroon pigmentation. These shifts are accompanied by the gain and later loss of delphinidin-derived anthocyanins, resulting in a return to biosynthetic pathways relying exclusively on cyanidin-based pigments (**Figure 2 & 3**). Such transitions are consistent with broader patterns observed across angiosperms, where evolutionary gains and losses of delphinidin production frequently underpin transitions between red, purple, and blue floral phenotypes (Wessinger and Rausher 2014; Noda 2018; Ogutcen et al. 2020; Roguz et al. 2020; Edwards et al. 2021; Wheeler et al. 2023, Stone & Wessinger, 2024).

While the biochemical basis of these floral colour transitions is predictable, the underlying genetic mechanism in blue-flowered Caladeniinae appears highly unusual. In most flowering plants, delphinidin production is typically mediated by F3′5′H enzymes belonging to the CYP75A clade, yet growing evidence indicates that this canonical pathway may be functionally compromised across Orchidaceae. CYP75A transcripts were frequently rare or absent in our dataset, while *Phalaenopsis* CYP75A-grouped proteins were incapable of anthocyanidin 3′- or 5′-hydroxylation and exhibited atypical substrate-binding orientations relative to functional F3′5′H enzymes (Liang et al. 2020). Genome-wide analyses further revealed that many orchid CYP75A proteins share structural features with these apparently non-functional proteins (Li et al. 2023), pointing towards broader functional erosion of CYP75A within the family.

Against this background, we uncovered a duplicated CYP75B lineage, CYP75B-D, associated almost exclusively with delphinidin-producing Caladeniinae species (**Figure 4**). CYP75B-D possesses the derived GLAL motif, including Ala at the critical eighth position of substrate recognition site 6, a substitution previously shown to confer de novo 5′-hydroxylation activity in CYP75B enzymes (Seitz et al. 2006; Seitz et al. 2007; Liu et al. 2020). Its origin was also accompanied by episodic positive selection, consistent with functional divergence following duplication. Blue-flowered Caladeniinae may therefore have regained access to delphinidin biosynthesis through duplication and neofunctionalisation of CYP75B-D rather than modification of a canonical CYP75A-mediated pathway. To our knowledge, this represents the first evidence for recruitment of F3′5′H-like activity by a CYP75B lineage within Orchidaceae, and potentially within monocots more broadly.

The co-option of duplicated CYP75B genes for 5′-hydroxylation has so far been documented in Asteraceae, where independently evolved CYP75B-derived F3′5′H activities contribute to blue and bluish-purple floral pigmentation in Asteroideae, Cichorioideae and Carduoideae (Seitz et al. 2007). The Caladeniinae therefore appear to represent an independent recruitment of this unusual mechanism within Orchidaceae. Related CYP75B-mediated routes may underpin other rare delphinidin-rich blue orchids, including *Vanda coerulea* and *V. coerulescens*, within the predominantly cyanidin-pigmented Epidendroideae (Tatsuzawa et al. 2004; Iwashina 2015). CYP75B-D may thus represent a key evolutionary innovation that enabled access to an otherwise rarely occupied region of orchid floral colour space.

Subsequent reversions to maroon pigmentation are most parsimoniously explained by loss of CYP75B-D expression or function rather than reversal of the D-type DFR state. The persistence of D-type DFR in maroon-flowered *C. ustulata* indicates that disruption of CYP75B-D may be sufficient to restore a cyanidin-only pathway despite retention of the derived DFR state (**Figure 2 & 5**; **Supplementary Data S5**). Potential mechanisms include regulatory silencing, disruptive coding mutations or complete loss of the duplicated gene. Transposable element-mediated changes are particularly plausible in Orchidaceae, where abundant copia and gypsy LTR retrotransposons occur throughout intergenic and intragenic regions and have been implicated in genome expansion and floral trait evolution (Trávníček et al. 2019; Chumová et al. 2021; Li et al. 2022; Russo et al. 2024; Li et al. 2025; Zhang et al. 2026). Genome-scale comparisons between blue- and independently derived, maroon-flowered lineages will be required to distinguish among these mechanisms.

### A shift in DFR substrate specificity preceded the evolution of blue floral pigmentation in Caladeniinae

In addition to CYP75B-D, the emergence of D-type DFR provides insight into floral colour diversification across the Caladeniinae. Although all blue-flowered taxa carried the derived D-type DFR, its occurrence in non-blue taxa indicates that the N-to-D transition pre-dated blue floral pigmentation (**Figure 5**). This is notable because monocots are dominated by N-type DFRs, whereas D-type variants are rare and frequently associated with delphinidin-rich systems (Johnson et al. 2001; Marin-Recinos and Pucker 2024; Smith and Rausher 2011; Wheeler et al. 2022). Unlike CYP75B-D, D-type DFR was not associated with gene duplication or strong evidence of adaptive protein evolution, suggesting that this functional transition involved limited sequence change within an otherwise conserved protein background.

Single amino acid substitutions within the conserved 26-amino-acid DFR substrate-binding region can substantially alter substrate preference and catalytic efficiency (Johnson et al. 2001; Petit et al. 2007; Haselmair-Gosch et al. 2018; Mei et al. 2019). In *Camellia sinensis*, conversion from N-type to D-type DFR reduced activity towards dihydroquercetin (DHQ) while increasing catalytic efficiency towards dihydromyricetin (DHM) (Mei et al. 2019); conversely, replacing Asp with Asn in *Petunia × hybrida* restored DHQ activity to levels comparable with DHM (Haselmair-Gosch et al. 2018). D-type DFR may therefore have represented an enabling evolutionary change that favoured reduction of the delphinidin precursor DHM. Previously documented in Orchidaceae only from *Glossodia major* (Wong et al. 2024), this derived state extends across the blue-flowered Caladeniinae and may have facilitated the subsequent recruitment of CYP75B-D.

Coordinated changes in DFR and flavonoid hydroxylase activity have accompanied floral colour transitions in several angiosperm systems. Transitions from blue delphinidin- to red pelargonidin-based pigmentation in *Iochroma* and *Penstemon*, and to cyanidin-rich phenotypes in *Phlox*, involve structural and regulatory changes affecting CYP75B and CYP75A activity (Hopkins and Rausher 2011; Smith and Rausher 2011; Wessinger and Rausher 2014; Wessinger and Rausher 2015; Wheeler et al. 2021). In *Lysimachia arvensis*, blue flowers similarly exhibit elevated expression of D-type DFR and F3′5′H, whereas orange flowers show reduced F3′5′H and increased N-type DFR expression (Sánchez-Cabrera et al. 2021). Together, these observations support a stepwise model in which altered DFR substrate preference preceded CYP75B-D neofunctionalisation, with their subsequent combination enabling efficient delphinidin biosynthesis and blue floral pigmentation in the Caladeniinae.

### Coordinated reorganisation of anthocyanin pathway expression accompanied blue floral pigmentation

Blue-flowered Caladeniinae exhibited a highly integrated anthocyanin biosynthetic pathway (ABP) expression architecture, contrasting with the partitioned organisation of maroon-flowered lineages (**Figure 6**; **Supplementary Figure S2**). The strong contribution of CYP75B-D to transcriptomic differentiation, together with its integration within the blue-flowered co-expression network, provides independent expression-level support for its association with delphinidin biosynthesis.

The negative associations of *FLS* with *ANS* and *arGST* provide further insight into the organisation of this network. Competition between FLS and the anthocyanin branch for shared dihydroflavonol substrates is well established (Luo et al. 2016; Yuan et al. 2016), potentially explaining the opposing relationship between *FLS* and *ANS*. The FLS–arGST association may reflect an additional interaction between these pathways, as quercetin, a flavonol produced downstream of FLS, can inhibit arGST activity affecting anthocyanin biosynthesis and sequestration (Morette et al. 2026). Conversely, flavonols can enhance blue pigmentation through co-pigmentation with delphinidin-derived anthocyanins, inducing bathochromic shifts and stabilising anthocyanin chromophores (Yoshida et al. 2009; Houghton et al. 2021). Blue pigmentation may therefore depend on a finely balanced relationship among anthocyanin biosynthesis, flavonol production, and co-pigmentation. Comparable co-regulated modules accompany floral colour loss and blue-to-red transitions across Solanaceae (Larter et al. 2018; Larter et al. 2019; Wheeler et al. 2023).

Anthocyanin pigmentation is largely controlled by MYB-bHLH-WD40 (MBW) transcription factor complexes (Albert et al. 2014). Classical studies in *Arabidopsis thaliana* distinguished early and late biosynthetic genes regulated primarily by SG7 and SG6/SG5 R2R3-MYBs, respectively (Xu et al. 2015). Higher-order mutant analyses nevertheless indicate that SG6 and SG5 MYBs can also activate early biosynthetic genes, allowing coordinated regulation across broader flavonoid modules (Sielmann et al. 2026). Similar complexity occurs in orchids: SG5 R2R3-MYBs regulate *F3H*, *DFR* and *ANS* in *Phalaenopsis* and *Gymnadenia* (Hsu et al. 2015; Kellenberger et al. 2019), while SG6 MYBs in *Dendrobium nobile* influence both anthocyanin and flavonol biosynthesis (Wang et al. 2026).

Although structural mutations are central to many floral colour transitions, regulatory divergence is also a major driver of pigment evolution (Wessinger and Rausher 2014; Wheeler et al. 2021; Wheeler et al. 2022). The contrasting expression architectures observed here suggest that blue floral pigmentation in the Caladeniinae involved pathway-wide regulatory change alongside the evolution of CYP75B-D and D-type DFR. Identifying SG6 anthocyanin regulators, SG7 flavonol regulators and SG4 repressors, together with their bHLH and WDR partners, will help resolve this regulatory architecture. MADS-box transcription factors also influence orchid floral pigmentation (Wang et al. 2026), raising the possibility that interactions between developmental and pigmentation regulators contributed to the gain and loss of blue flowers.

### Floral colour and pollinator shifts in Caladeniinae coincide with paleoclimatic events

Increasing evidence suggests that floral colour evolution is shaped by interactions between pollinator communities and environmental conditions across Australian ecosystems (Gascoigne et al. 1948; Caruso et al. 2019; Dalrymple et al. 2020; Sullivan and Koski 2021). The Caladeniinae provide a compelling example: blue-flowered lineages emerged c. 18–14 Ma (O’Donnell et al. 2024), spanning the Middle Miocene Climate Optimum (MMCO, 18–14 Ma) and the subsequent Middle Miocene Climatic Transition (MMCT) around 14 Ma (Byrne et al. 2011; Crisp and Cook 2013) (**Figure 7**). Global cooling and increasing Australian aridification during the MMCT drove major ecological restructuring and shifts in floristic composition (Crisp and Cook 2007; Lewis et al. 2008; Byrne et al. 2011). The divergence of the Western Australian *Cyanicula* s.s. and *Elythranthera* from the eastern Australian *Glossodia* c. 13 Ma (O’Donnell et al. 2024) also coincides with formation of the arid Nullarbor Plain c. 13–14 Ma, consistent with numerous east-west disjunctions across the Australian flora (Crisp and Cook 2007). These temporal associations suggest that climatic change helped shape the diversification and biogeography of blue-flowered Caladeniinae.

**Figure 7.**
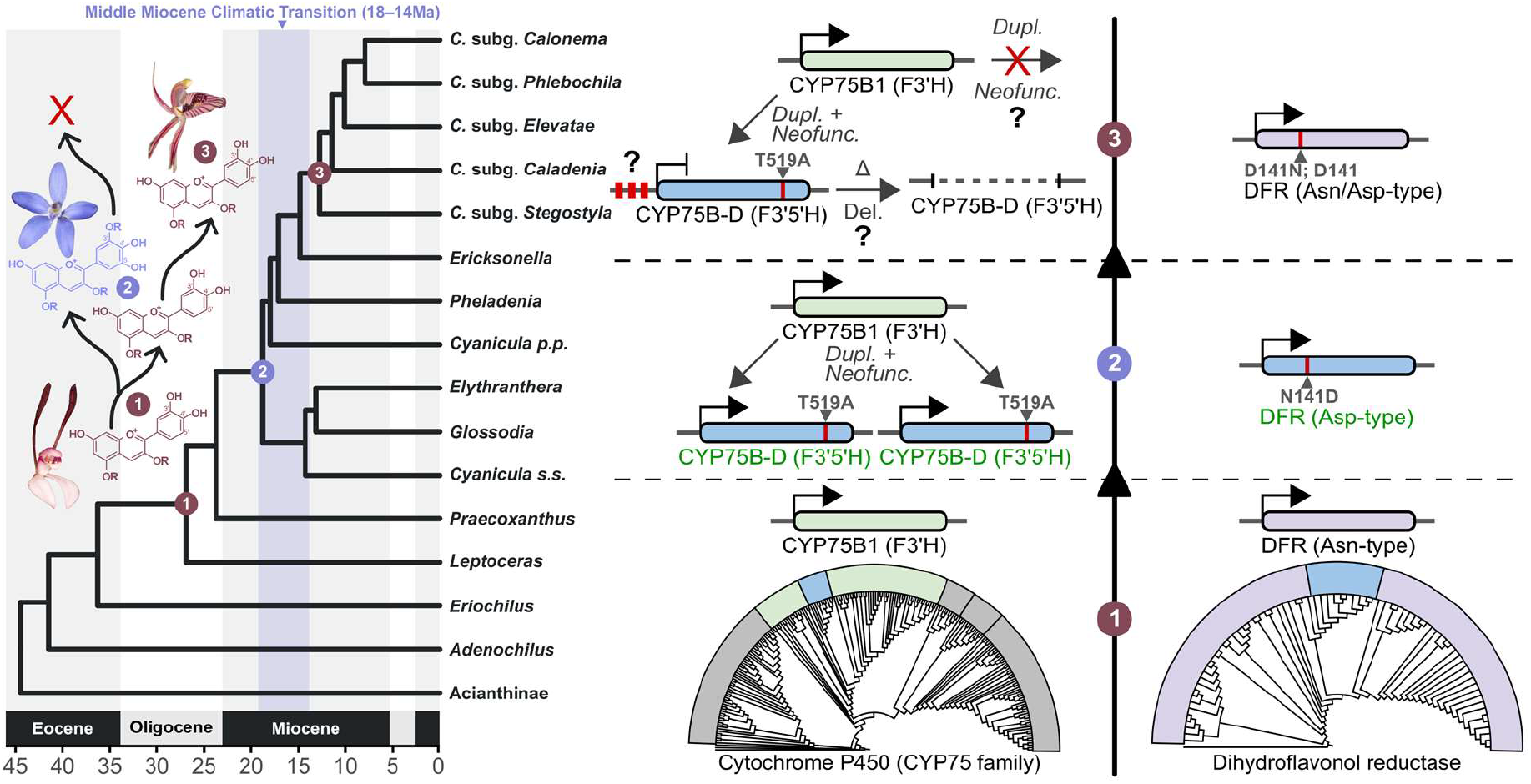
Conceptual model of floral colour evolution in the Caladeniinae. Timetree adapted from O’Donnell et al. (2024) illustrating the inferred timing of major anthocyanin biosynthetic pathway innovations associated with transitions in floral pigmentation and pollination strategy within the subtribe (transitions 1-3). Arrows indicate inferred gene duplication and neofunctionalization events, whereas crossed red arrows denote the absence of these innovations. Key amino acid transitions associated with floral colour evolution are shown at relevant nodes, including the CYP75B SRS6 and DFR substrate-binding site shifts. Question marks (?) indicate hypothesised silencing of CYP75B-D (red promoter region) or loss of the duplicated gene (Δ) during reversions from blue- to maroon-flowered lineages. Associated anthocyanin chemistry is shown for each transition, with red and blue anthocyanin structures representing cyanidin- and delphinidin-based pigmentation, respectively. The timing of these transitions is presented in the context of major palaeoenvironmental events and pollinator shifts discussed in the text. Abbreviations: **CYP75B**, flavonoid 3′-hydroxylase; **CYP75B-D**, duplicated CYP75B lineage associated with the apparent recruitment of 5′-hydroxylase activity; **DFR**, dihydroflavonol 4-reductase.

The gain and subsequent loss of delphinidin-based blue pigmentation also coincided with shifts in pollination strategy. Visual signals are particularly important in bee-pollinated systems, whereas sexually deceptive orchids rely strongly on olfactory mimicry and pollinator-specific semiochemicals (Peakall 2023). The emergence of blue-flowered lineages may therefore reflect a broader shift in pollinator attraction strategies during the MMCO–MMCT interval. The molecular innovations identified here provide a mechanistic basis for this change in floral signalling, although causal links among climate, pollinator transitions and floral colour remain to be tested.

Subsequent reversions to maroon pigmentation accompanied transitions to sexual deception of thynnine wasps. However, thynnine-based sexual deception arose considerably earlier in the distantly related Drakaeinae *sensu stricto*, c. 29.9–24.6 Ma (Weston et al. 2014; O’Donnell et al. 2024), indicating that diversification of these orchid lineages was not tightly coupled to the initial emergence of this pollination niche. Instead, repeated transitions to thynnine pollination may have imposed comparable selective pressures on floral traits, including colour patterning and visual contrast (Gaskett et al. 2017; Basist et al. 2021).

Similar colour-pollinator associations occur elsewhere in Diurideae. Within Thelymitrinae, blue-flowered, bee- and beetle-pollinated *Epiblema* and *Thelymitra* contrast with maroon-flowered *Calochilus*, several species of which are pollinated by sexually deceive male wasps (Jones and Gray 1974; Bower and Branwhite 1993; Bernhardt and Edens-Meier 2014; Weston et al. 2014). The *Thelymitra-Calochilus* divergence occurred at approximately the same time as the emergence of blue-flowered Caladeniinae, around 14 Ma (O’Donnell et al. 2024). A comparable association is evident at the subtribe scale between the sexually deceptive Cryptostylidinae and blue-flowered Thelymitrinae (Weston et al. 2014; O’Donnell et al. 2024). These recurrent associations suggest that similar ecological pressures may have favoured convergent combinations of floral colour and pollination strategy across Australian orchids.

## Conclusions

The transition from ancestral maroon to derived blue flowers, followed by reversions to maroon pigmentation, represents one of the most striking evolutionary patterns within the Caladeniinae. By integrating phylotranscriptomics, anthocyanin profiling, molecular evolution and gene expression analyses, we show that this transition was accompanied by coordinated changes in pathway structure, protein function and expression architecture. Most notably, we identified a duplicated CYP75B lineage strongly associated with delphinidin-producing taxa, together with a derived D-type DFR that appears to have established an enabling biochemical background for blue floral pigmentation. The emergence of blue flowers was further accompanied by substantial reorganisation of anthocyanin pathway expression, which coincided with pollinator transitions during a period of major palaeoenvironmental change. Together, these findings suggest that blue floral pigmentation in the Caladeniinae arose through the interaction of molecular innovation, regulatory evolution, pollinator-mediated selection and ecological opportunity.

## Acknowledgements

We acknowledge Australia’s First Nations Peoples as the Traditional Custodians of the land and waters on which this work was conducted, recognizing their historical and ongoing stewardship of its genetic resources, and pay our respects to all Elders past and present. We thank the staff at the ANU Joint Mass Spectrometry Facility for assistance with LC–MS operation, Noushka Reiter and Richard Dimon from the Royal Botanical Garden Victoria for tissue sampling, and Eran Pichersky for sequencing some early ancillary transcriptomes utilized in this study.

## Data availability statement

The transcriptome assemblies presented in this study have been deposited in NCBI Transcriptome Shotgun Assembly (TSA) Database and under the NCBI BioProject (https://www.ncbi.nlm.nih.gov/bioproject/) accession PRJNAXXXXXX.

